# Assessing cats’ (*Felis catus*) sensitivity to human pointing gestures

**DOI:** 10.1101/2022.03.12.484069

**Authors:** Margaret Mäses, Claudia A.F. Wascher

## Abstract

A wide range of non-human animal species have been shown to be able to respond to human referential signals, such as pointing gestures. The aim of the present study was to replicate previous findings showing cats to be sensitive to human pointing cues (Miklósi et al. 2005). In our study, we presented two types of human pointing gestures - momentary ipsilateral (direct pointing) and momentary cross-body pointing. We tested nine rescue cats in a two-way object choice task. On a group level, the success rate of cats was 74.4 percent. Cats performed significantly above chance level in both the ipsilateral and cross-body pointing condition. Trial number, rewarded side and type of gesture did not significantly affect the cats’ performances in the experiment. On an individual level, 5 out of 7 cats who completed 20 trials, performed significantly above chance level. Two cats only completed 10 trials. One of them succeeded in 8, the other in 6 of these. The results of our study replicate previous findings of cats being responsive to human ipsilateral pointing cues and add additional knowledge about their ability to follow cross-body pointing cues. Our results highlight that a domestic species, socialised in a group setting, may possess heterospecific communication skills. Further research is needed to exclude alternative parsimonious explanations, such as local and stimulus enhancement.

## Introduction

A wide range of non-human animal species has been shown to be able to respond to human referential signals, such as pointing gestures (Krause et al., 2018; Miklósi & Soproni, 2006). Pointing presents a species-specific human communicative signal (Bard et al., 2021). The ability of humans to understand pointing with a hand as an object-directed action develops at the age of 9 to 12 months (Woodward & Guajardo, 2002). The development of pointing comprehension in humans and non-human animals is likely a result of learning, social experience and interactions (Miklósi & Soproni, 2006). A variety of non-domesticated mammalian taxa, including dolphins (*Tursiops truncatus*; Herman et al., 1999), elephants (*Loxodonta africana*; Smet & Byrne, 2013), bats (*Pteropus*; Hall et al., 2011) and sea lions (*Zalophus californianus*; Malassis & Delfour, 2015), have demonstrated following some form of human pointing. Several studies have examined the understanding of human pointing cues in chimpanzees (*Pan troglodytes*) and other great apes, specifically in the object choice task. Initial studies suggested subjects were relatively unsuccessful (Kirchhofer et al., 2012; Povinelli et al., 1997). However, recent studies suggest systematic confounds rather than differences between species to cause this effect (Clark et al., 2019; Clark & Leavens, 2019; Hopkins et al., 2013). For example the rearing environment affects the performances of apes in pointing tasks and individuals reared in complex environments outperformed individuals reared under standard conditions (Lyn et al., 2010; Russell et al., 2011).

When it comes to domestic animals, goats (*Capra hircus*; Kaminski et al., 2005; Nawroth et al., 2020), pigs (*Sus scrofa domestica*; Nawroth et al., 2016), horses (*Equus caballus*; Proops et al., 2010), cats (*Felis catus*; Miklósi, et al., 2005), and most prominently, dogs (*Canis familiaris*; Bhattacharjee et al., 2020; Bräuer et al., 2006; Hare et al., 1998; Soproni et al., 2002; Tauzin et al., 2015) have been shown to follow pointing signals. In the case of dogs (in particular, the domestication process has been considered to have shaped the evolution of their remarkable socio-cognitive skills that allow them to successfully communicate with humans (Hare et al., 2002). However, this hypothesis is challenged by a range of wild canids such as wolves (*Canis lupus*), coyotes (*Canis latrans*), and foxes (*Vulpes vulpes*) responding to human pointing gestures, as well as socialisation with humans affecting dogs’ performance, with pet dogs outperforming dogs housed in kennels and shelters (reviewed in Krause et al., 2018).

Despite domestic cats being one of the most popular pets and very well adapted to human environments, their cognition has been studied notably less than that of dogs (Shreve & Udell, 2015). In a previous study, Miklósi, et al. (2005) demonstrated cats’ abilities to follow human pointing were comparable to the abilities of dogs doing so, whereas they performed more poorly compared to dogs in attention-getting behaviour. In another study however, cats responded to the attentional state of a person when presented with an unsolvable task (Zhang et al., 2021). Cats are able to use human gaze as a referential signal (Pongrácz et al., 2019). Performance of cats has recently also been tested in other cognitive tasks. For example they have been shown to differentiate between different quantities (Pisa & Agrillo, 2009), they are able to mentally represent the location of non-visible objects (Takagi et al., 2021) and reproduce a human’s familiar action on an object (touch it with hand/paw or face) (Fugazza et al., 2021).

Nevertheless, the body of research on socio-cognitive capacities of cats remains currently considerably small. Interestingly, it has been suggested that the process of cat domestication is different from that of other domestic species, as it was driven by a mutualistic relationship with humans and was subject to significantly less strict artificial selection (Clutton-Brock, 1994; Serpell, 2013). Cat domestication can even be claimed to have been self-initiated (Driscoll et al., 2009). Another aspect worth taking into account is that, compared to most other species studied in the context of social cognition, cats have an arguably less social lifestyle, as their ancestors were primarily solitary (Bradshaw, 2016). One might expect that these evolutionary peculiarities have a negative effect on cats’ responsiveness to human communicative signals.

One of the measures by which referential cues can be categorized is their duration, the signal being either momentary or dynamic (Miklósi & Soproni, 2006). For momentary pointing, the signaller keeps the arm in the pointing position for only a second (Miklósi et al., 2005). On the other hand, when giving a dynamic cue, the signal is terminated after the receiver has responded (Miklósi & Soproni, 2006). The momentary cues are arguably more similar to naturally occurring communicative interactions than dynamic cues, as the subject has to remember the signal before making a choice. In the present study, we aimed to test whether cats follow the human momentary ipsilateral (direct) pointing cues in a two-way choice task, choosing the target indicated with the referential signal at above-chance level, thus replicating the findings of Miklósi et al. (2005). Additionally, we tested whether cats follow the human momentary cross-body pointing cues in a two-way choice task. As the cross-body form of the signal was most likely novel to the subjects, we expected the cats to be more successful in following ipsilateral pointing cues. If cats show the ability to respond accurately to different forms of pointing cues this could be indicative of an ability to generalize and potentially referential understanding.

## Methods

### Ethical considerations

The present study received ethical approval from the School Research Ethics Panel of Anglia Ruskin University. The study was approved by and conducted at Pesaleidja cat shelter in the Republic of Estonia. This study complies with the national regulations on ethics and research on animals in Estonia.

### Standards for openness and transparency

We report how we determined our sample size, all data exclusions, all manipulations, and all measures in the study.

#### Study subjects

The experiment was conducted during summer 2020 (29 June - 12 August). Study subjects were housed in a rehoming centre in Tallinn, managed by Pesaleidja NGO. A total of approximately 200 cats were roaming free in different indoor spaces (10 – 51 m^2^; 0.5 cats per a square metre; Jaroš, 2018), nine of which participated in the study. Cats were individually tested in a separate room.

The cat’s suitability to participate in the experiment was evaluated in three stages (similar to the method of Miklósi et al. (2005), with certain alterations described below). Firstly, the potential subject was approached by the experimenter (M.M.), who sat down next to the individual, and petted it for one minute. If the cat did not leave during this time or express fearful behaviour (e.g. flattened ears (Bennett et al., 2017; Deputte et al., 2021; Gourkow et al., 2014); whiskers held against face; dilated pupils; becoming immobile/freezing; piloerection; arched back; tensely crouched body position; tail tucked tightly between the legs or around the body; hissing or other agonistic vocalizations (Tavernier et al., 2020)); of any kind, the experimenter guided the subject into the testing room (5.5 m^2^), either by allowing it to follow the experimenter or alternatively carrying it for a maximum of ten seconds. After separation the subject was given time to explore the testing room. Here the subject was isolated from its conspecifics for the duration of the experiment, the doors were closed to prevent the other cats from entering. With those individuals not initially comfortable, i.e. expressing fearful or stressed behaviour (e.g. attempting to hide (Bennett et al., 2017; Gourkow et al., 2014); yowling (Tavernier et al., 2020) and standing fixated to one of the closed doors; pacing back and forth (Gourkow et al., 2014)), with the novel setting, the experimenter sat on the floor and petted them, calmly allowing them to walk around, as well as offering some food. If the cat continued showing signs of stress after five minutes, the experimenter allowed it to exit the room and excluded it from any further testing. As a last stage of habituation, the experimenter put some food into one of the test bowls (green silicone muffin cases) and introduced it to the cat by allowing it to smell the bowl. We used small amounts of wet cat food, as recommended by the shelter staff, as a reward throughout the experiment. Rewards were given to the subjects in addition to their normal diet. The bowl was then placed on the floor, approximately one metre from the subject. The cat was allowed to approach it and eat the food. If the cat was motivated to approach the bowl and showed interest in eating the food, it passed the third stage and was included in the final experiment. This stage additionally familiarised the cats with the bowl containing a food reward. Twenty cats passed the first stage of preliminary testing, but some of them did not habituate to the novelty of testing room environment quickly enough, were not food motivated or showed a persistent side bias (description below). Consequently, ten subjects participated in the final experiment. However, we decided to exclude one of them from data analyses due to side bias. The remaining nine subjects all completed a minimum of ten experimental trials. Seven of them completed 20 trials.

#### Study design

As the cats’ everyday diet was provided to them *ad libitum*, timing of the experiment was not dependent on the feeding regime. Before every trial and out of site of the subject, approximately the same amount of food, positioned as similarly as possible, was put into both test bowls (paying attention to prevent visual and odour-induced bias of choice). Next, a bit of food liquid was smeared onto the inner walls of a third silicone bowl, serving as ‘bait’ distracting the cats while the experimenter got into position. The subject was attracted to a position approximately two metres away the experimenter’s final position. The experimenter simultaneously placed the test bowls in front of them, the middle line between the bowls at an approximate distance of 0.5 metres. The experimenter then made an attention-drawing sound (common utterance used for calling cats in the local area: ‘ks-ks’) and presented the pointing cue when the subject was looking in the direction of the experimenter.

We tested cats’ responses to ipsilateral pointing to the left (IL), with the left arm and index finger pointing at the container on the left side of the experimenter, ipsilateral pointing to the right (IR), with the right arm and index finger pointing at the container on the right side of the experimenter, cross-body pointing to the left (CL), with the right arm and index finger pointing at the container on the left side of the experimenter, and cross-body pointing to the right (CR), with the left arm and index finger pointing at the container on the right side of the experimenter. The experimenter maintained a neutral body posture and gaze direction, at all times, while performing the pointing gestures. After pointing, the subject could choose one of the bowls. The cat was considered to make a choice when it looked into the bowl or reached into it with its paw. When the choice corresponded to the direction of the gesture, the cat was allowed to eat the reward from the ‘correct’ bowl. When the choice was ‘unsuccessful’, both bowls were picked up before the subject was able to eat the food. In the case of the subject not making a choice (*e.g.*, walked between the test bowls and straight to the experimenter or walked away), the experimenter repositioned themselves and repeated the trial. In one subject, the experimenter could not lead the subject to refocus, and therefore, stopped the session and continued on another day. Order of trials in the four conditions (IL, IR, CL, CR) was pseudo-randomized. Each condition was presented five times in a total of 20 test trials. Each condition was not repeated more than twice in a row and the type or direction a maximum of three times.

If the subject continuously chose the bowl on the same side for four consecutive trials, regardless of the signal, we considered this as an indication for the subject developing a side bias. In this case, the positioning of the experiment was switched to the opposite side of the room, which seemed to be effective with four subjects. One subject, who had passed the three stages of preliminary testing but reached for the bowl on the right side for ten consecutive trials, was excluded from further participation in the experiment.

#### Data analyses

Data was analysed by M.M., indicating correct, *i.e.* the cat chose the side which was pointed towards, and incorrect, *i.e.* the cat chose the side which was not pointed towards, responses. An inter-observer reliability analysis was conducted on 30 % of randomly chosen trials, which were coded by a second observer (C.A.F.W.). Inter-observer agreement was 100 %. Statistical analyses were performed in R 4.0.3 (The R Foundation for Statistical Computing, Vienna, Austria, http://www.r-project.org). A generalised linear mixed model (GLMM) with a binomial distribution and logit link was used to investigate differences in performance between different conditions in the package lme4 (Bates et al., 2015). Trial outcome (successful or unsuccessful) was the response variable, the signal type (ipsilateral or cross-body pointing), location (left or right) and the trial number (1-20) were included as fixed factors, and the subject identity as a random effect. To assess multicollinearity between fixed factors, we calculated variance inflation factors (VIFs) using the vif function in the package car (Fox & Weisberg, 2011). VIFs for all factors were below 2, indicating that there was no issue with multicollinearity (Zuur et al., 2009). To describe the variance explained by our models, we provided marginal and conditional R^2^ values that range from 0 to 1 and described the proportion of variance explained by the fixed effects and by the fixed and random effects combined, respectively (Nakagawa & Schielzeth, 2013). We calculated marginal and conditional R^2^ values using the r.squaredGLMM function in MuMIn (version 1.15.6; Bartoń, 2019). We conducted exact, two-tailed binomial tests to investigate whether cats used pointing gestures significantly above chance. Cohen’s h (h) was calculated as a measure of effect size, using the package pwr (Champely, 2020). In individuals who completed the full 20 trials we further conducted binomial tests to see whether individuals were successful above chance level. All datasets and the R^2^ script used to conduct the statistical analyses are available as supplementary files.

## Results

On a group level, the success rate of cats was 74.4 %. Cats performed significantly above chance level in both the ipsilateral pointing (Binomial test: p<0.001, h = 1.287, [95% confidence intervals = 0.702 - 0.884]) and cross-body pointing condition (Binomial test: p = 0.002, h = 0.823, [95% confidence intervals = 0.564 - 0.78]; Figure 1). Trial number (GLMM: estimate ± standard deviation = −0.009 ± 0.032, z-value = −0.284, p = 0.776), rewarded side (GLMM: estimate ± standard deviation = 0.238 ± 0.371, z-value = 0.372, p = 0.709) and type of gesture (GLMM: estimate ± standard deviation = 0.667 ± 0.374, z-value = 1.78, p = 0.074) did not significantly affect the cats’ performances in the experiment (intercept: GLMM: estimate ± standard deviation = 0.797 ± 0.473, z-value = 1.685, p = 0.091). Overall, 2 % of the variation in performance was explained by all fixed factors together (R^2^ marginal), and an additional 2 % of the variation in performance was explained by the random factor (individual, R^2^ conditional). On an individual level, 5 out of 7 cats who completed 20 trials, performed significantly above chance level (individual 2: Binomial test: p = 0.011, h = 1.287, [95% confidence intervals = 0.563 - 0.942], individual 3: Binomial test: p < 0.001, h = 1.854, [95% confidence intervals = 0.683 - 0.987], individual 4: Binomial test: p = 0.503, h = 0.402, [95% confidence intervals = 0.36 - 0.808], individual 5: Binomial test: p = 0.011, h = 1.287, [95% confidence intervals = 0.563 - 0.942], individual 6: Binomial test: p = 0.041, h = 1.047, [95% confidence intervals = 0.508 - 0.913], individual 7: Binomial test: p = 0.823, h = 0.2, [95% confidence intervals = 0.315 - 0.769], individual 8: Binomial test: p = 0.002, h = 1.55, [95% confidence intervals = 0.621 - 0.967]; Figure 2). Two cats only completed 10 trials. One of them succeeded in 8, the other in 6 of these.

**Figure 1.**
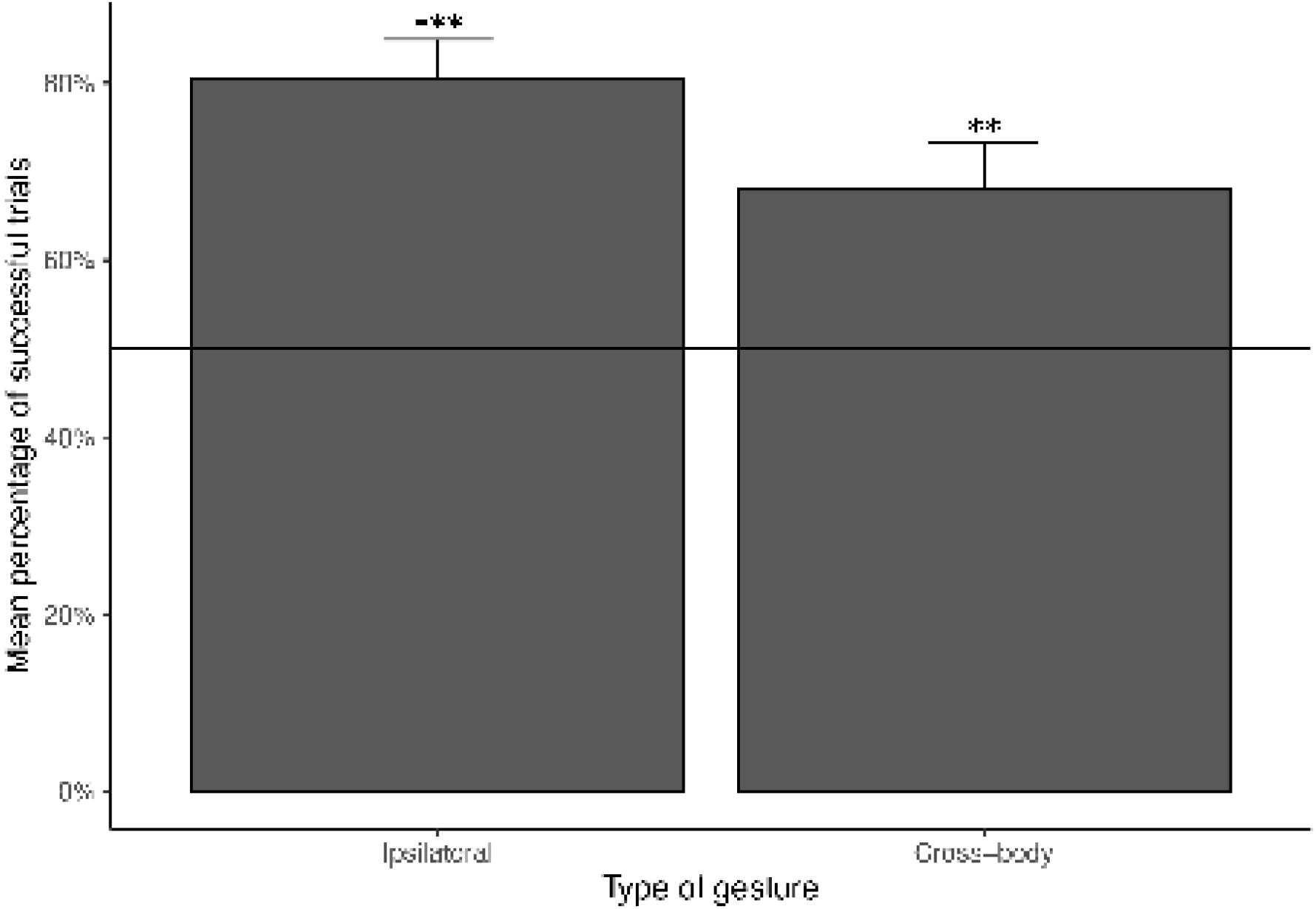
Mean percentage of trials plus standard error where the cats followed ipsilateral pointing and cross-body pointing. Full line represents 50 % chance level. **P < 0.01; ***P < 0.001.

**Figure 2:**
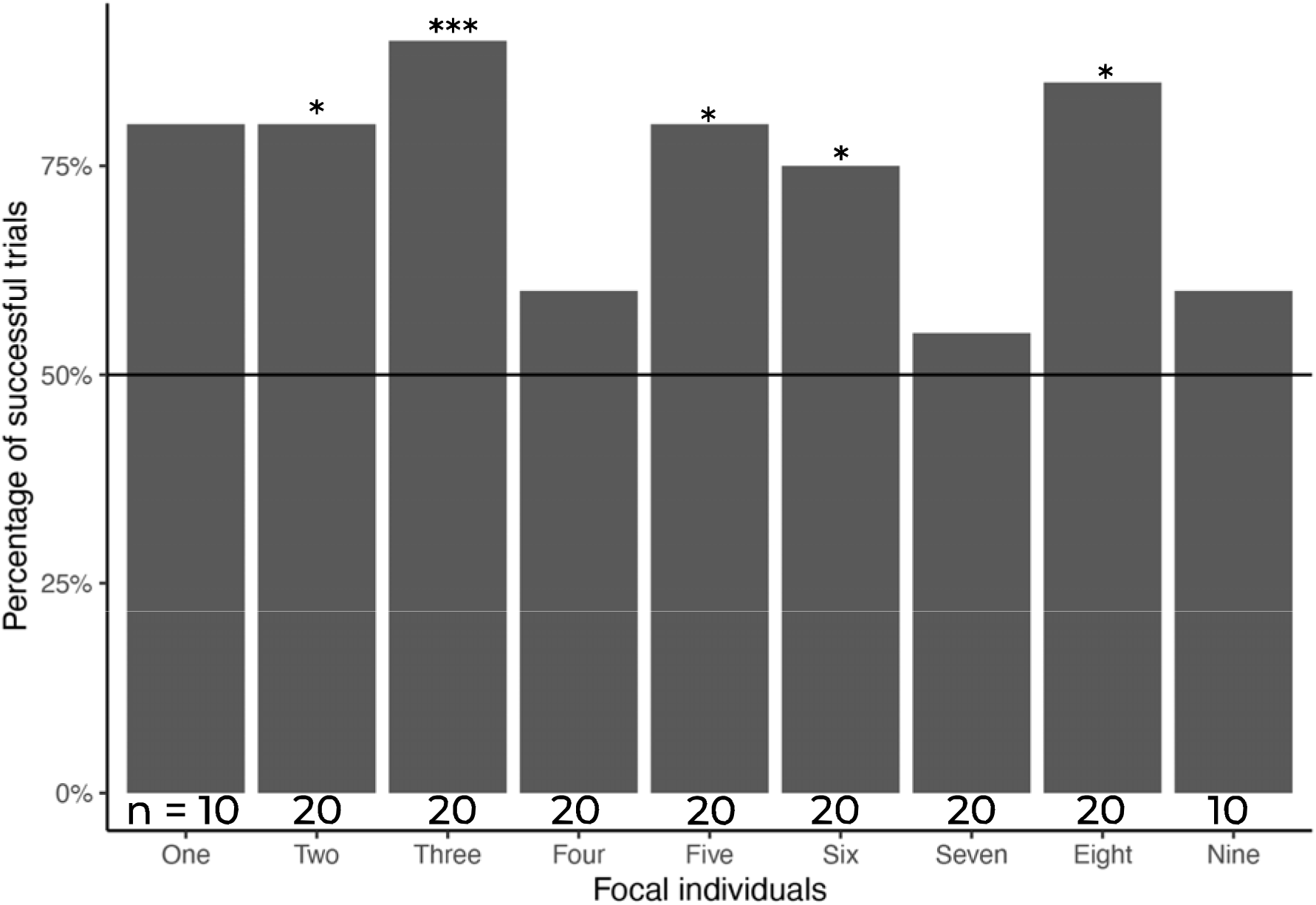
Percentage of successful trials for each focal individual. Sample size (n) indicates the number of trials per individual. Full line represents 50 % chance level. Binomial test: *P < 0.05; ***P < 0.001.

## Discussion

The results of the present study show the cat’s ability to follow human ipsilateral pointing gestures (Bard et al., 2021), which replicates findings of a previous study (Miklósi *et al.* 2005). Additionally, we show cats are sensitive to cross-body pointing cues. Out of the seven individuals tested in 20 trials, five followed human pointing cues significantly more often than expected by chance. We did not find a significant difference in performance between ipsilateral pointing and cross-body pointing. The ability to follow human cross-body pointing gestures has been previously shown in a wide variety of species (for a review see Pack, 2019).

Our results show that, similarly to dogs and some other species, the more solitary cats use communicative cues from humans. Cognitively, different mechanisms could be involved in the ability of cats to follow human communicative cues, such as stimulus or local enhancement as well as cue learning. If the subjects’ choices had been influenced by rapid learning, the performance would be expected to improve over the testing trials (Kaminski et al., 2005; Malassis & Delfour, 2015; Miklósi et al., 2005). The trial number showed no significant influence on trial outcome. Thus, we conclude that subjects are not learning to follow a point during the course of testing.

From an evolutionary perspective, the finding that cats are sensitive to human pointing cues is interesting, as cats and their ancestors do not normally experience conspecifics pointing. It has previously been suggested that the process of domestication has selected for socio-cognitive abilities that enable domesticated species to better communicate with humans compared to wild species (Hare et al., 2002). In a previous study Miklósi et al. (2005) directly compared dogs’ and cats’ abilities to follow human pointing cues and attention-getting behaviour. While dogs and cats did not differ in their ability to follow human pointing cues, cats lacked some components of attention-getting behaviour compared with dogs, in accord with the domestication hypothesis. However, recent studies directly comparing human-socialized dogs and wolves show the wolves to outperform dogs, in contrast to the domestication hypothesis (Range & Marshall-Pescini, 2022; Udell et al., 2008, 2010). To investigate the effects of domestication on cats’ performance, it would be necessary to conduct comparable assessments of the sensitivity to human pointing gestures in socialized individuals of wild cats (*Felis lybica* and/or *Felis silvestris*; Pongrácz, Szapu & Faragó, 2019).

Importantly, our study adds to a growing body of literature highlighting that less social species are able to master socio-cognitive tasks. For example, non-social reptiles (*Geochelone carbonaria*) and fish (*Spinachia spinachia*; *Cottus gobio*; *Barbatula barbatula*; *Platichthys flesus*) have been shown to use social information (Webster & Laland, 2017; Wilkinson et al., 2010). It has been previously suggested that socialisation with humans can enable animals to acquire communicative skills which allow them to respond to cues from heterospecifics (Kaminski et al., 2005; Nawroth et al., 2020; Proops et al., 2010; Range & Marshall-Pescini, 2022). However, we would like to highlight that there are more parsimonious alternative explanations, namely following human pointing via local and stimulus enhancement, which in the present experiment cannot be ruled out.

Compared to similar studies with cats or dogs, where the experiments have been conducted in the owners’ homes (*e.g.*, Miklósi et al., 2005; Pongrácz et al., 2019), the standardisation of the testing environment in the current study could be considered an advantage. In a previous study, family-owned dogs outperformed kennel-housed dogs in their capacity to understand human pointing gestures (D’Aniello et al., 2017; Lazarowski & Dorman, 2015). As mentioned above, cats do not use pointing cues in conspecific communication, hence any previous experience of the cats that participated in this study with pointing must have come from human-cat interactions in the shelter or before cats came to the shelter. The shelter environment also means that cats have been living in a group situation for extended periods of time, which could have allowed them to acquire certain socio-cognitive skills that are less evident in cats without this extensive social experience with conspecifics.

Similar to all other studies on animal cognition and behaviour, we need to consider potential sample bias of our study population as outlined in the STRANGE framework (Webster & Rutz, 2020). We must consider the social background of focal subjects and as mentioned above, we acknowledge previous experience with conspecifics and heterospecifics (humans) in the group-housed cats. Self-selection could have affected our results, as from the 200 cats in the shelter, we only tested nine individuals who voluntarily participated in the experiment, based on the cat being comfortable when isolated from the group and interacting readily with the human experimenter. It could very well be that this procedure excluded focal subjects who are less responsive to human pointing cues. Future investigations into individual differences in performance and cats’ abilities to follow human pointing cues would be desirable. As our focal subjects are shelter cats, we have very little information about their rearing history and past experience, and no information about their genetic make-up. Moreover, our experiment was of a short-term nature, capturing the cats’ responses during a short-term period. We did not intend to investigate potential natural changes in responsiveness, *e.g.*, seasonal or ontogenetic changes, and these areas should be considered for future studies.

## Acknowledgements

We thank Johanna Miedel, the manager of Pesaleidja, for granting permission to work with the cats. We would also like to thank the rest of the team of staff and volunteers at Pesaleidja shelter for their guidance and cooperation on site. We also thank two anonymous reviewers and the editor, Prof. Dorothy Munkenbeck Fragaszy for their comments, which greatly improved the article.

## Author contributions

Conceptualization: MM and CAFW; Methodology: MM and CAFW; Investigation: MM; Formal analysis: MM and CAFW; Supervision: CAFW; Writing: MM and CAFW.

